## Supplementary Tables for "Molecular and electrophysiological features of spinocerebellar ataxia type seven in induced pluripotent stem cells"

| **S Table 1: Primary antibodies** | | | | |
| --- | --- | --- | --- | --- |
| **Antibody** | **Full name** | **Species** | **Supplier and catalogue number** | **Dilution** |
| AFP | Alpha-fetoprotein | Mouse | Abcam, ab54745 | 1:100 |
| ASA | Anti-sarcomeric alpha actinin | Rabbit | Abcam, ab137346 | 1:100 |
| ATXN7 | Ataxin 7 | Rabbit | Thermo Scientific, PA1749 | 1:400 |
| CRX | Cone-rod homeobox | Sheep | R&D Systems, AF7085 | 1:20 |
| FOXA2 | Forkhead box A2 | Rabbit | Abcam, ab23630 | 1:1000 |
| GABA | γ-aminobutyric acid | Rabbit | Sigma-Aldrich,  A2052 | 1:500 |
| GFAP | Glial fibrillary acidic protein | Rabbit | Abcam, ab7260 | 1:100 |
| NES | Nestin | Mouse | Abcam, ab6320 | 1:1000 |
| NP | Nucleocapsid protein of Sendai virus | Mouse | Gift from Mahito Nakanishi | 1:1500 |
| OCT4 | POU class 5 homeobox 1 | Rabbit | R&D systems, KQX0409011 | 1:200 |
| RCVRN | Recoverin | Rabbit | Millipore, ab5585 | 1:1000 |
| SMA | Smooth muscle actin | Mouse | Abcam, ab7817 | 1:100 |
| TRA-1-60 (PODXL) Alexa-488 conjugated | Podocalyxin like | Mouse | Millipore, MAB4360A4 | 1:200 |
| βIII-Tubulin | Tubulin, beta 3 class III | Mouse | Abcam, ab78078 | 1:300 |

| **S Table 2: Secondary antibodies** | | | | |
| --- | --- | --- | --- | --- |
| **Antibody** | **Dye** | **Species** | **Supplier** | **Dilution** |
| Anti-sheep | Alexa-488 | Donkey | Jackson Immunoresearch  **713-545-003** | 1:500 |
| Anti-rabbit | Cy3 | Donkey | Jackson Immunoresearch  **711-165-152** | 1:500 |
| Anti-mouse | Alexa-488 | Goat | Jackson Immunoresearch  **115-545-003** | 1:500 |

| **S Table 3: Primer sequences** | | |
| --- | --- | --- |
| **Gene** | **Primer** | **Sequence (5' to 3')*** |
| ATXN7 | Atxn7 CAG RNA F | HEX-CGAGCTTTCAGAATGCAGC |
|  | Atxn7 CAG RNA F | CACTTCAGGACTGGGCAGAG |
| B-ACTIN | FORWARD | Proprietary Primer Design UK |
|  | REVERSE | Proprietary Primer Design UK |
| NANOG | FORWARD | CAGCCCCGATTCTTCCACCAG |
|  | REVERSE | CGGAAGATTCCCAGTCGGGTT |
| SOX2 | FORWARD | GGGAAATGGGAGGGGTGCAAA |
|  | REVERSE | TTGCGTGAGTGTGGATGGGAT |
| OCT3/4 | FORWARD | GACAGGGGGAGGGGAGGAGC |
|  | REVERSE | CTTCCCTCCAACCAGTTGCCC |
| MITF | FORWARD | TTCACGAGCGTCCTGTATGCAGAT |
|  | REVERSE | TTGCAAAGCAGGATCCATCAAGCC |
| NRL | FORWARD | GGTCCTAGTCCCAGCTCTTC |
|  | REVERSE | TCGTCCAATCCACATGAGAATTA |
| OTX2 | FORWARD | TGCAGGGGTTCTTCTGTGAT |
|  | REVERSE | AGGGTCAGAGCAATTGACCA |
| PAX6 | FORWARD | CGGAGTGAATCAGCTCGGTG |
|  | REVERSE | CCGCTTATACTGGGCTATTTTGC |
| RCVRN | FORWARD | CCAGAGCATCTACGCCAAGT |
|  | REVERSE | CACGTCGTAGAGGGAGAAGG |
| RHO | FORWARD | GTCGATTCCACACGAGCACTG |
|  | REVERSE | CCTCTCTGAATGGATACTTCGTC |
| RPE65 | FORWARD | GCCCTCCTGCACAAGTTTGACTTT |
|  | REVERSE | AGTTGGTCTCTGTGCAAGCGTAGT |
| ARR3 | FORWARD | TCACTTCCAAGTCATCACGG |
|  | REVERSE | GGTGTTGTCCTGGTTGATCC |
| GNAT1 | FORWARD | TAGCTGAGGGGAGTGCAAAT |
|  | REVERSE | CCTCAAAGACTGTGGCCTCT |
| ATXN7 | FORWARD | GCCAGCCGTGAACAATGTC |
|  | REVERSE | TTCCTCCCCGTGCTATTTTCA |
| BEX1 | FORWARD | GGAGGAGACTACAAGGATAGG |
|  | REVERSE | TCCTTTTCTTCATTTTCTTGGTT |
| DNAJA1 | FORWARD | AAAGGAGGAGAACAGGCAATTAA |
|  | REVERSE | TAGGGTTACTGAGAGCTGATGT |
| GRIA2 | FORWARD | CTATGGCATCGCAACACCTAA |
|  | REVERSE | GTCCTTGGCTCCACATTCAC |
| HSP27 | FORWARD | ACGAGCTGACGGTCAAGAC |
|  | REVERSE | GGGGGCAGCGTGTATTTCC |
| HSP70 | FORWARD | ATGGAATCTATAAGCAGGATCT |
|  | REVERSE | CACATACAGAAACTTGATAAGC |
| HSP105 | FORWARD | CCCGTCAGTCATATCATTTGGA |
|  | REVERSE | AATCTTTTGAAGTTAGACACCGTATT |
| OLIG1 | FORWARD | GTTTGGAGAGCTGTATTTAAGACT |
|  | REVERSE | TTCTAAGAAACCCCCAGGATTTA |
| UCHL1 | FORWARD | TGAAGCAGACCATTGGGAAT |
|  | REVERSE | TGTTTCAGAACTGATCCATCCT |
| PLCB3 | FORWARD | CCTTGGAAATCTTTGAGCGGTTC |
|  | REVERSE | ACTTCGTTGAGTCTCGGGTC |
